# Predicting seabird-vessel interactions from bird-borne data alone

**DOI:** 10.64898/2026.11.03.763666

**Authors:** Leia Navarro-Herrero, David March, Frederic Bartumeus, Jacob González-Solís

## Abstract

1. Seabirds often follow fishing vessels to scavenge discards and bait, benefiting from access to food but also facing the risk of bycatch. Exposure to bycatch is commonly assessed by analysing seabird-fishing interactions, typically through the spatiotemporal alignment of seabird and vessel tracking data. However, this approach may underestimate interactions when vessel tracking information is incomplete or unavailable.
2. We tested whether seabird-vessel interactions could be identified using seabird telemetry alone and whether fishing and nonfishing vessels could be distinguished. We analysed GPS, wet-dry, accelerometer and pressure data from 273 Scopoli’s shearwaters in the Northwest Mediterranean. We identified 779 vessel-attendance events using seabird GPS, Automatic Identification System (AIS) data and simulation-based tests to distinguish vessel attraction or following from chance encounters. We then trained random forest models to predict vessel-attendance from bird-borne movement and environmental metrics, testing different sensor combinations. Finally, we selected the GPS and environmental model for its performance and compatibility with GPS-only tracking data from 17 radar-equipped shearwaters, whose detectors identified nearby vessels independently of AIS.
3. Models predicting vessel-attendance from seabird movement metrics derived from GPS, wet-dry sensors and accelerometers achieved classification rates above 90% during training. However, when tested on new individuals, 42% of their positive predictions were false-positives. Models combining environmental metrics with either GPS or GPS and wet-dry data achieved classification rates around 75% during training and generalised consistently across individuals, maintaining classification rates of 70% or above. Attendance events involving fishing (n = 482) and nonfishing vessels (n = 297) could not be reliably distinguished.
4. Across 45 trips of radar-equipped shearwaters, the GPS and environmental model discriminated vessel-attendance in hourly bins against AIS detections (AUC = 0.85), supporting assessment of spatiotemporal attendance patterns. At the event level, only 17% of 227 predicted vessel-attendance events were supported by AIS or radar detections. However, radar detected nearby vessels during 19 predicted events lacking AIS support, highlighting the value of complementary vessel detection.
5. Policy implications. This study presents a vessel-independent framework for assessing seabird-vessel interactions based solely on bird-borne data, complementing fisheries monitoring programmes, helping to identify gaps in vessel-tracking systems across poorly monitored marine regions, and informing future assessments of bycatch risk.

## 1 INTRODUCTION

Seabirds are frequently observed following vessels at sea, often engaging in scavenging discards, baits, or offal (Sherley et al., 2019; Votier et al., 2010). Yet, consuming such predictable human-provided food sources instead of their natural prey can have adverse effects on the health and survival of these species (Furness, 2003; Lewison et al., 2004; Oro et al., 2013). For example, discards, which may be of poor nutritional quality, can result in reduced caloric intake in the diet of seabirds (Grémillet et al., 2008). Moreover, birds may accidentally collide with, become entangled in, or hook themselves on fishing gear (bycatch) while attempting to catch fish, which remains one of the most significant threats to seabird populations (Dias et al., 2019; Lewison et al., 2014; Žydelis et al., 2009). These impacts raise important conservation concerns and have driven increased interest in studying seabird-fishing interactions over the past three decades.

Combining seabird tracking with vessel tracking at intervals of a few minutes (1-10 minutes) has been crucial for assessing seabird-fishing interactions and bycatch risk (Clark et al., 2020; Corbeau, Collet, Orgeret, et al., 2021; Navarro-Herrero et al., 2025). However, the two main vessel tracking systems used, Vessel Monitoring System (VMS) and Automatic Identification System (AIS), have significant limitations. Regarding availability, VMS data are often held by governments, and high-resolution AIS data can incur high financial costs. Concerning reliability, VMS data may be updated only every 1-2 hours in Europe, while AIS, though more frequent (5-10 min), suffers from notable data gaps caused by reception issues, system saturation in high traffic areas, equipment failures, or intentional deactivation (Ford et al., 2018; Park et al., 2023; Shepperson et al., 2018; Welch et al., 2022; Paolo et al., 2024). Moreover, EU regulations (EC 2244/2003) exempt vessels <12 m (VMS) and <15 m (AIS), excluding much of the small-scale fishing fleet from bycatch assessments (Pott & Wiedenfeld, 2017).

Alternative approaches have been developed to detect vessels near seabirds when vessel tracking data are not available. One such approach uses bird-borne tags to pick up radar emissions from nearby vessels, which are installed for navigational safety (Weimerskirch et al., 2018). However, these tags do not reveal the identity of the vessels, and thus cannot distinguish fishing vessels, which pose a bycatch risk to seabirds, from nonfishing vessels such as passenger ships. In addition, radar tags may underestimate seabird-fishing interactions, either due to tag malfunction (Carneiro, Dias, et al., 2022; Weimerskirch et al., 2020), or the limited adoption of the radar systems, particularly among small vessels.

Another approach, which until recently remained largely conceptual, is to infer when a bird is following a vessel solely from how it moves (i.e., its movement patterns), as discussed by Corbeau et al. (2019). Initial empirical attempts to test this idea found that albatross behaviour associated with vessel-attendance was difficult to distinguish reliably from natural foraging patterns (Carneiro, Dias et al., 2022). More recently, however, two studies have reported promising results by predicting interactions between albatrosses and fishing vessels using movement variables derived from bird-borne sensors (Le Scornec et al., 2025; Rutter et al., 2025). Nevertheless, several limitations remain. First, both studies focused on albatrosses in regions dominated by industrial fisheries, and it therefore remains unknown whether this approach generalises to species with different movement ecology or to regions where recreational, passenger, or small-scale fishing vessels may also contribute substantially to interactions. Second, achieving robust predictive performance on independent datasets remains a key challenge that has not always been addressed. Le Scornec et al. (2025) showed that model performance decreased substantially when validated on independent individuals, raising questions about the transferability of such models across individuals and populations. Finally, both studies based their predictions on single-sensor data streams, despite earlier work exploring the integration of multiple sensor types (Carneiro, Dias et al., 2022). There is therefore still scope to investigate whether integrating multiple sensor types can capture a broader range of seabird behaviours and improve the robustness of movement-based interaction inference.

Recent advances in biologging miniaturization now enable monitoring of seabird behaviour using a single tag or a lightweight combination of tags that integrate multiple sensors. GPS tracking has long been the standard approach for studying seabird movement, providing fine-scale information on flight paths at the scale of minutes, even seconds. GPS tags can also be combined with additional bird-borne sensors that capture aspects of behaviour not detectable from location data alone and record activity at much finer temporal resolutions. For instance, tri-axial accelerometers measure acceleration using gravitational force to determine bird orientation, quantifying energy expenditure and flight behaviours such as flapping and gliding (Collins et al., 2015). Wet-dry sensors detect water contact to identify landings and sustained flights (Dehnhard et al., 2020; Dias et al., 2012), and barometric pressure sensors provide diving depth and duration (Browning et al., 2017).

In this study, we used a long-term biologging dataset (GPS, accelerometers, wet-dry, pressure, and radar detectors) from Scopoli’s shearwaters (*Calonectris diomedea*) in the Northwest Mediterranean Sea to investigate whether distinct seabird movement patterns occur when birds actively associate with fishing vessels. Unlike the albatrosses examined in previous studies with the same objective (Carneiro, Dias et al., 2022; Le Scornec et al., 2025; Rutter et al., 2025), this species frequently interacts not only with industrial fisheries but also extensively with small-scale fisheries from the region, posing a substantial threat due to high bycatch risk (Cortés et al., 2017; Cortés & González-Solís, 2018; Genovart et al., 2017; Reyes-González et al., 2021; Soriano-Redondo et al., 2016). Moreover, these waters are also one of the busiest maritime regions worldwide, with intense shipping, cruise tourism and recreational boating (March et al., 2021a), further increasing potential interactions between seabirds and nonfishing vessels (Navarro-Herrero et al., 2024; March et al., 2026).

Here, we used GPS trajectories of Scopoli’s shearwaters with vessel AIS data to identify vessel-attendance events beyond chance co-ocurrences, following March et al. (2026). We then tested whether movement metrics alone (e.g. speed, energy expenditure, or diving behaviour) could predict (a) whether a seabird was interacting with a vessel, and (b) whether the interaction involved a fishing or nonfishing vessel. Given the expected difficulty of predicting vessel interactions from movement metrics alone, we also evaluated whether adding environmental metrics associated with each GPS position, such as distance to the coast, distance to the colony, and time of day, could enhance predictive accuracy (Granadeiro et al., 2011; Orben et al., 2021; Reyes-González et al., 2021; Torres et al., 2011). Finally, we applied the selected model to complete foraging trips and used bird-borne radar detections as an independent source of nearby vessel presence, including vessels not represented in AIS data. This allowed us to assess the potential and limitations of inferring vessel-attendance from bird-borne data when vessel-tracking information is incomplete.

## 2 MATERIAL AND METHODS

### 2.1 Data collection

#### 2.1.1 Seabird telemetry data

Between 2015 and 2021, we tagged Scopoli’s shearwater individuals that were breeding or prospecting at Cala Morell colony (Balearic Islands, Spain, Northwest Mediterranean Sea; Fig. 1A) with three types of GPS tags: GPS-only tags (weight ∼18 g, Perthold Engineering LLC), GPS combined with tri-axial accelerometers and pressure sensors (weight ∼17 g, Technosmart Europe S.r.l.) or GPS combined with radar detectors (weight ∼19 g, Sextant Technology Ltd).

**Figure 1.**
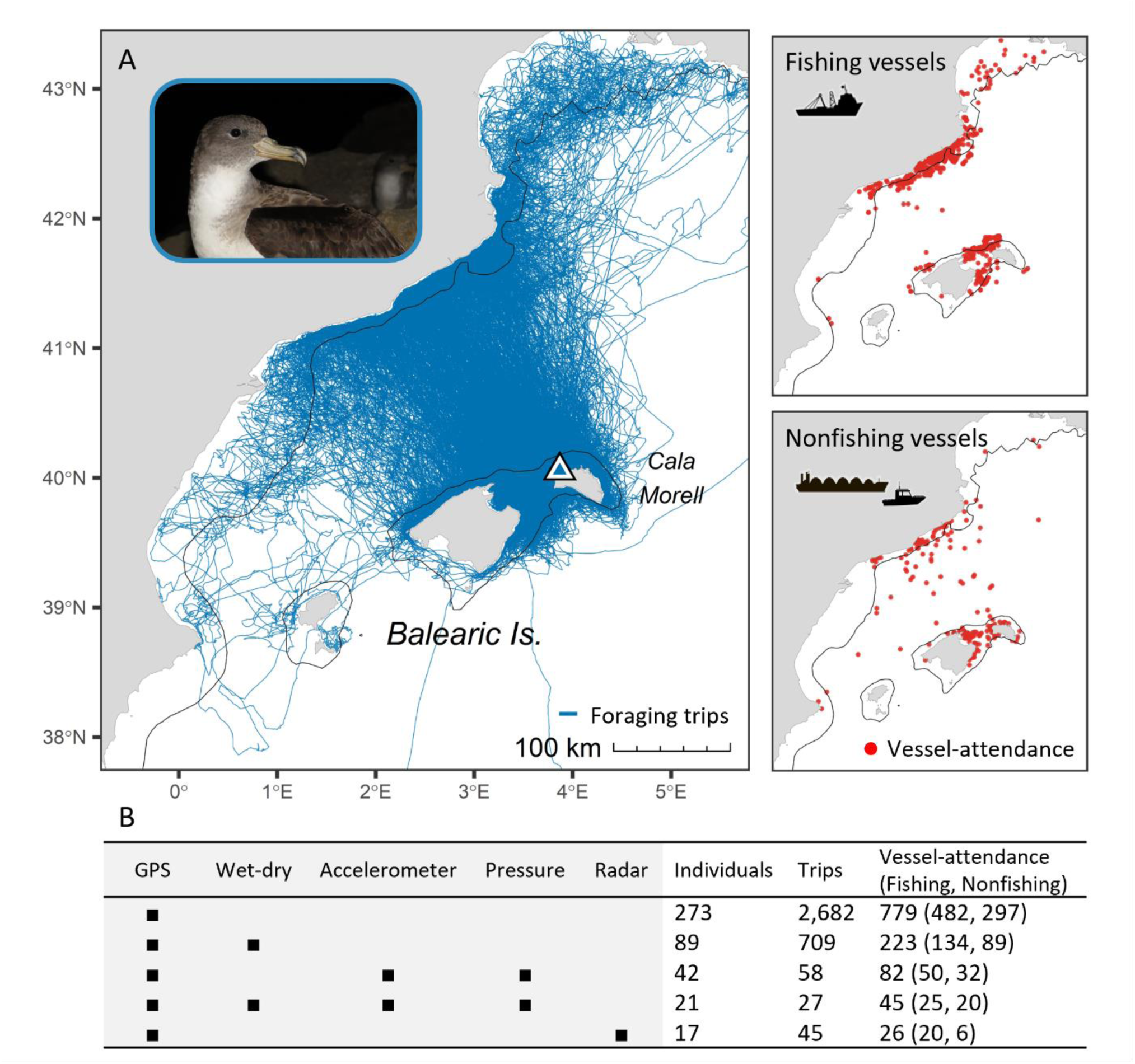
Bird-borne data collected from Scopoli’s shearwaters. (A) Foraging trips (blue lines) and seabird– vessel interactions (red dots). Vessel-attendance events were identified by applying simulation-based inference to threshold-defined association events derived from seabird GPS and vessel AIS data. Vessel-attendance segments comprised each classified event and the preceding 30 min. The triangle indicates the breeding colony, and the black line the 200 m isobath. Photograph by Montserrat Vanerio. (B) Sensor combinations used in this study and their respective sample sizes. Sample sizes are not additive, as individuals may be represented in more than one sensor category.

The GPS tags were attached to the dorsal feathers using waterproof Tesa tape. The GPS tags, when combined with the accelerometers, were consistently deployed to account for the bird orientation along the X (head-tail), Y (right-left), and Z (dorsoventral) axes. Occasionally, individuals were tagged simultaneously with a wet-dry sensor tied to a plastic ring attached to the leg (weight ∼3 g, GLS C-250 Migrate Technology). The total weight of the tags for simultaneous tagging remained at <4% of the weight of the bird (∼600 g), thus avoiding potential disruptions of seabird foraging behaviour (Passos et al., 2010). Wet-dry sensors were retrieved concurrently with the GPS tags after approximately five days of deployment or at the end of the fieldwork season. GPS was set up to record fixes every 5 minutes. We removed fixes with unrealistic flight speeds (above 80 km h^-1^) with the *SDLfilter* R package (Shimada et al., 2012). For each individual, GPS positions were classified using the Expectation-Maximisation binary Clustering algorithm (EMbC; Garriga et al., 2016), based on speed and turning angle. The four resulting states (intensive search, extensive search, resting and relocating) were reduced to three by grouping both search states as foraging (Reyes-González et al., 2021).

The accelerometer and pressure sensor were set up following the recommendations for the same species (Cianchetti-Benedetti et al., 2017). The accelerometer measured the vector sum of gravity (i.e., static acceleration, a constant at 1 g ≈ 9.8 m s^-2^ at the earth surface) and animal movement (i.e., dynamic tag acceleration). It was set up to record tri-axial accelerometer data within a sensitivity range of −8 to 8 g every 0.04 seconds (25 Hz). Pressure sensor was set up to record pressure in millibars (mBar, with an accuracy of 5 mBar ≈ 0.05 m) every second. Wet-dry sensor was set up to record wet or dry states every 6 seconds.

Finally, a radar detector, which scans marine radar emissions in a radius of ∼5 km (based on Weimerskirch et al., 2018), was set up to scan for 2 minutes every 5 minutes, following a previous study on the same species (Navarro-Herrero et al., 2024).

Individuals were handled following relevant regulations and Association for the Study of Animal Behaviour (ASAB) and the Animal Behavior Society (ABS) Guidelines for the Use of Animals in Research, with all permits approved by the *Conselleria de Medi Ambient, Agricultura i Pesca* (permits: CEP24/2015, ANE06/2015, CEP30/2016, ANE08/2016, ANE24/2017, ANE27/2018, CEP21/2019, ANE19/2020, ANE18/2021).

#### 2.1.2 Vessel tracking data

Vessel tracking trajectories (Terrestrial AIS data) were obtained from the Balearic Islands Coastal Observing and Forecasting System (March et al., 2021a). AIS messages provided location fixes every 5 minutes, the Maritime Mobile Service Identity (MMSI) code, and vessel type. We removed AIS messages that were duplicated or had invalid MMSI codes (i.e., codes without nine digits or not starting with digits 2-7, which identify individual vessels and exclude other AIS transmitter types). We classified vessels into fishing and nonfishing vessels (i.e., tanker, cargo, passenger or recreational vessels) using AIS metadata (March et al., 2021). In addition, because vessel-type information in AIS messages is self-reported and may contain errors or inconsistencies, fishing vessels were identified as those listed in the Global Fishing Watch (https://globalfishingwatch.org) or the European Fishing Fleet Register (https://webgate.ec.europa.eu/fleet-europa).

### 2.2 Identifying vessel-attendance segments

We identified when seabirds intentionally approached vessels following the framework developed by March et al. (2026). We first split seabird GPS tracks into foraging trips (consecutive at-sea locations between colony departure and return) using the *track2KBA* R package (Beal et al., 2021). We then linearly interpolated seabird and vessel trajectories to 5 minute intervals using the *move* R package (Kranstauber et al., 2018) to obtain simultaneous locations (Chisholm et al., 2019). Interpolated positions were retained only if the nearest observed vessel fix occurred within 5 minutes.

We defined association events as periods during which a seabird remained within 1.5 km of a vessel for at least 15 minutes, allowing interruptions of up to 30 minutes within the same event (March et al. 2026). Because such proximity may reflect either a directed behavioural response or incidental co-occurrence, each association was evaluated independently for attraction and following behaviour using 100 simulated null trajectories. Attraction was assessed by testing whether the association was unlikely to arise under seabird movement independent of vessel position, whereas following was assessed by testing whether the observed association persisted longer than expected under the null model. Associations classified to be significant as attraction and/or following at p < 0.05 were hereafter termed vessel-attendance events. Thus, association denotes threshold-defined spatiotemporal co-occurrence, whereas vessel attendance denotes an association supported by significant attraction and/or following. Full details of the simulation procedure and sensitivity analyses are provided in March et al. (2026).

For each vessel-attendance event, we extended the underlying association event by including the 30 minutes immediately preceding the start of the vessel-attendance event accounting for the possibility that seabirds may begin altering their movement in response to vessels before coming into close proximity (Corbeau et al., 2019; Votier et al., 2010; March et al. 2026). The resulting trajectory, comprising this pre- association period and the complete vessel-attendance event, was termed as vessel-attendance segment. Vessel-attendance segments therefore varied in duration according to the duration of each attendance event. Instances in which a seabird simultaneously attended multiple fishing vessels were excluded to ensure that each vessel-attendance event could be confidently assigned to a specific vessel.

### 2.3. Predictive modelling of vessel attendance

Our first objective was to assess whether seabird movement patterns differed between vessel-attendance and natural-movement (i.e. periods with no detected vessel-attendance). Our second specific objective was to determine whether seabird movement patterns during vessel-attendance events could distinguish fishing from nonfishing vessels.

#### 2.3.1. Response variable and trajectory segmentation

We used Random Forest classification models (RF; Breiman, 2001) using a binary response variable tailored to each objective. For first objective, the response variable distinguished between vessel-attendance (1) and natural-movement (0). The model unit was either a vessel-attendance segment a or natural-movement segment. Natural-movement segments were represented by pseudo-absences generated by randomly selecting trajectory segments where no vessel interaction was detected from AIS data. Since not all vessels have AIS or broadcast their positions, some pseudo-absences might still include unseen vessel interactions, which could add noise to this dataset. Selected natural-movement segments had characteristics comparable to vessel-attendance segments. Specifically, for each vessel-attendance segment, we selected a natural-movement segment matching its total duration and predominant EmbC-derived behavioural state. The predominant state of each segment was determined from the proportion of time spent in each state. Further details on the procedure used to generate pseudo-absence segments are provided in Appendix S1.

For second objective, the response variable classified only vessel-attendance segments, distinguishing associations with fishing vessels (1, N = 482) from those with nonfishing vessels (0, N = 297).

#### 2.3.2. Predictor variables

Both objectives used the same set of predictor variables. For each model unit (vessel-attendance segment or natural movement segment), we calculated a range of movement and behavioural metrics commonly used in movement ecology (Williams et al., 2019) from the available sensors. GPS-derived metrics (N = 15) described distance travelled and displacement, step length, path sinuosity and straightness, speed, and time spent in each EMbC-derived behavioural states (Appendix S2). Wet-dry metrics (N = 7) described landing frequency, transitions between wet and dry states, and the duration and proportion of time spent in each state (Appendix S3). Accelerometer metrics (N = 28) described acceleration along the three body axes (surge, sway and heave), body orientation (pitch and roll), and overall dynamic body acceleration (ODBA and VeDBA), including their mean and variability within segments (Appendices S4-S5). Pressure metrics (N = 7) described diving activity, including dive frequency, maximum depth, and the duration and phases of dives (Appendices S4 and S6). Finally, environmental metrics derived from GPS data (N = 4) described time of day through solar position (altitude and azimuth) and the bird’s distance to the coast and breeding colony (Appendix S7).

#### 2.3.3. Model fitting and performance

For both classification objectives, we built Random Forest (RF) models for the four sensor combinations from bird-borne sensor data: (1) GPS (N = 273 individuals); (2) GPS with wet-dry sensor (N = 89 individuals); (3) GPS with accelerometer and pressure sensors (N = 42 individuals); and (4) GPS with accelerometer, pressure, and wet-dry sensors (N = 21 individuals; Fig. 1B). Accelerometer and pressure sensors were integrated into the same tags and were therefore always analysed together. Sensor combinations were cumulative, such that individuals with multiple sensors could contribute data to more than one model. Training and testing subsets were generated independently for each sensor combination, with approximately 80% of observations allocated to training and the remaining 20% to testing. Observations from each individual were assigned entirely to either the training or the testing dataset, ensuring that no bird contributed to both subsets and allowing model evaluation on previously unseen individuals.

The number of predictor variables was first reduced using the minimally biased variable selection algorithm implemented in MUVR2 (Shi et al., 2019). Variable selection was performed using the training dataset to avoid information leakage into the independent testing dataset. RF models were fitted using the *ranger* and *caret* R packages (Kuhn, 2008; Wright & Ziegler, 2017), with 1,000 trees and the Gini splitting rule. Hyperparameters were tuned by varying the number of variables randomly sampled at each split (mtry, from 1 to the total number of predictors) and the minimum number of observations required in terminal nodes (min.node.size).

Depending on the number of individuals available for each model, cross-validation was conducted using leave-one-out, five-fold or ten-fold approaches. Because trajectory segments from the same individual are not independent, we applied grouped cross-validation, assigning all observations from each individual to the same fold (Roberts et al. 2017; March et al. 2021b). The optimal hyperparameter combination was selected by maximising Youden’s J statistic (sensitivity + specificity − 1) during cross-validation, which balances the correct detection of positive cases with the avoidance of false detections (i.e. identifying vessel-attendance segments while avoiding misclassifying natural-movement in the first objective, and identifying fishing vessel-attendance while avoiding misclassifying nonfishing vessels in the second objective). The optimal hyperparameter combination identified during cross-validation was used to fit the final RF model, which was subsequently evaluated on the independent testing dataset. Variable importance was quantified using the impurity-based importance measure.

Building on previous studies addressing the same prediction problem (Carneiro, Dias et al., 2022; Rutter et al., 2025; Le Scornec et al., 2025), we evaluated model performance using sensitivity (the proportion of true positives correctly identified) and specificity (the proportion of true negatives correctly identified), calculated separately for the training and independent testing datasets.

### 2.4. Predicting vessel attendance along entire foraging trips

We aimed to apply an RF model to predict vessel-attendance events across complete foraging trips. We anticipated that some predictions would not coincide with AIS-derived vessel-attendance. Some of these discrepancies may result from the inherent classification error of the model, whereas others may represent actual vessel-attendance events involving vessels not broadcasting AIS. To assess whether these apparently unsupported predictions coincided with nearby vessels, we required a model that could also be applied to trips where birds carried radar detectors (N = 45 trips, Fig. 1B), which detect nearby vessels regardless of AIS transmission. Because radar detectors were rarely deployed alongside additional sensors owing to weight constraints, only models based on GPS-derived data were suitable for this assessment. Among these, we selected the model incorporating GPS-derived movement and environmental variables because it showed better predictive performance than the model based on GPS-derived movement metrics alone.

Because RF models were trained on trajectory segments rather than individual GPS locations, complete foraging trips were also split into segments before making predictions. To represent vessel-attendance events of different durations, we segmented each complete trip three times, using segment lengths representing short (45 minutes), intermediate (80 minutes) and long events (135 minutes, see Appendix S8 for details on segment classification). We then applied the RF model to all segments from the three segmentations. The predicted probability for each segment was assigned to all GPS locations within that segment. Because the same GPS location could occur in segments of different lengths, it could receive more than one predicted probability. We therefore averaged these probabilities to obtain a single prediction for each GPS location.

Because consecutive GPS locations were temporally autocorrelated, we grouped locations into consecutive 1-hour bins to integrate predictions from the three segmentations and evaluate model performance while reducing temporal dependence. This temporal scale was consistent with previous work on the same species (Navarro-Herrero et al., 2024) and was supported by an autocorrelation analysis (Appendix S9).

We used the area under the curve (AUC) for this assessment using the *dismo* R package (Hijmans, 2017), comparing a predicted probability of vessel-attendance (ranging from 0 to 1) with an observed binary state (0 = no vessel attendance; 1 = vessel attendance) for each bin. For each bin, the predicted probability was calculated by averaging the location-level vessel-attendance probabilities, while the observed state was assigned a value of 1 when the bin contained at least one AIS-derived vessel-attendance location, and 0 otherwise. Because vessel-attendance bins were substantially less frequent than non-attendance bins, AUC was estimated from 500 balanced resamples, each containing all observed vessel-attendance bins and an equal number of randomly selected observed non-attendance bins. We report the mean AUC and its 95% confidence interval across resamples, with 0.5 indicating no discrimination and 1 indicating perfect discrimination.

We next compared the temporal agreement between predicted vessel-attendance, AIS-derived vessel-attendance and radar detections within each individual. While AIS-derived attendance was represented as a binary state (0 = no vessel-attendance, 1 = vessel-attendance) and radar detections as vessel presence or absence (0 = no radar detection, 1 = radar detection), model predictions were probabilities of vessel-attendance ranging from 0 to 1. Predicted probabilities were therefore converted into binary vessel-attendance states using the probability threshold that maximised the sum of sensitivity and specificity (*spec_sens* criterion in the *dismo* R package). Consecutive detections within each data source (predictions, AIS and radar) separated by less than 30 min were then grouped into events. Isolated radar detections were excluded because they indicated only transient vessel presence. Finally, events were summarised by their start and end times and considered concordant when their temporal intervals overlapped within individuals.

A curated, user-friendly workflow containing the R code and a subset of the data required to reproduce the main analyses is provided as Supplementary Material accompanying this preprint.

## 3 RESULTS

### 3.1 Vessel attendance

Of the 273 tagged seabirds in this study, 66% (N = 180 individuals) attended a vessel at some point during their foraging trips, and these associations lasted on average 38.3 ± 33.8 min (mean ± SD). However, seabirds generally attended vessels more than once per trip (2.2 ± 2.2 times, mean ± SD), and these interactions accounted for 2.5 ± 3.2% (mean ± SD) of the total trip duration (5.6 ± 3.6 days each trip, mean ± SD). During vessel-attendance segments, seabirds spent more time foraging or relocating than resting (Appendix S10), and most often attended a single vessel at a time (∼90% of the vessel-attendances). We excluded 46 instances of simultaneous attendance at multiple fishing vessels. Overall, we identified 779 vessel-attendance segments, of which 62% (N = 482) involved fishing vessels (Fig. 1A).

### 3.2 Discrimination of vessel attendance

After applying variable selection with the MUVR algorithm, models classifying vessel-attendance from natural-movement retained only a subset of the original predictor variables. Between four and 12 of the 15 GPS metrics were retained, together with one of the seven wet-dry metrics, seven to eight of the 28 accelerometer metrics, and none of the seven pressure-derived metrics (Appendix S11).

Among all models evaluated, the model combining GPS (4 metrics), wet-dry (1 metric), and accelerometer (7 metrics) predictors achieved the highest performance in the training dataset, with both sensitivity (correct classification of vessel-attendance segments) and specificity (correct classification of natural-movement segments) above 90% (Table 1). GPS and accelerometer-derived metrics were among the most important predictors (Fig. 2A1). The four most important metrics showed that the vessel-attendance time window used to train the models was associated with faster movement, greater distances travelled (length and maximum length), and higher overall body movement (VeDBA) compared with natural-movement (Fig. 2B1). However, this performance was not maintained in new individuals, with specificity falling below 60% in the independent testing dataset, meaning that more than 40% of natural-movement segments were incorrectly classified as vessel-attendance (Table 1).

**Figure 2.**
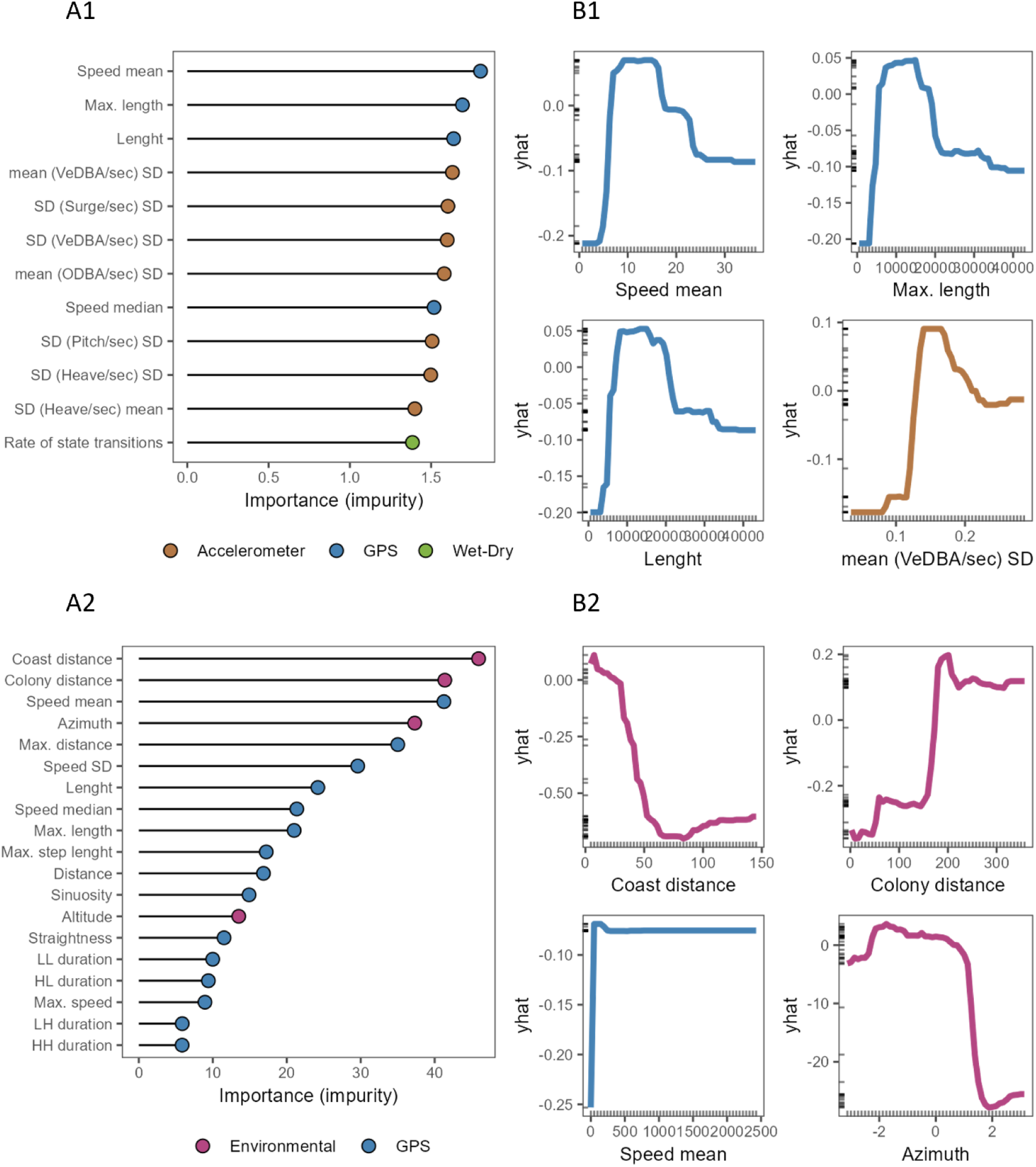
Variable importance (A1–A2) and partial dependence plots for the four most important predictors (B1–B2) of Random Forest models discriminating vessel attendance from natural movement. A1–B1 show the GPS, wet-dry and accelerometer model, which performed best during training; A2–B2 show the GPS and environmental model, which performed more consistently across training and testing and was subsequently applied to complete foraging trips. The ‘yhat’ denotes the model predicted values of the response variable. See Appendix S2-S5 for description of all variables.

**Table 1.** Performance metrics of Random Forest models discriminating vessel-attendance from natural-movement. Squares indicate the predictor groups included in each model. Models are ordered according to Youden’s J statistic. N indicates the number of trajectory segments in the training and testing datasets.

|  |  |  |  | Training |  |  |  | Testing |  |  |  |
|---|---|---|---|---|---|---|---|---|---|---|---|
| GPS | Wet-dry | Accel. | Envir. | Youden's J | Sensitivity | Specificity | N | Youden's J | Sensitivity | Specificity | N |
| ■ | ■ | ■ |  | 0.83 ± 0.23 | 0.93 ± 0.12 | 0.90 ± 0.14 | 62 | 0.44 | 0.86 | 0.58 | 33 |
| ■ | ■ | ■ | ■ | 0.81 ± 0.19 | 0.93 ± 0.12 | 0.88 ± 0.11 | 62 | 0.44 | 0.86 | 0.58 | 33 |
| ■ |  | ■ | ■ | 0.63 ± 0.18 | 0.90 ± 0.12 | 0.73 ± 0.20 | 140 | 0.43 | 0.83 | 0.60 | 27 |
| ■ |  | ■ |  | 0.60 ± 0.24 | 0.88 ± 0.17 | 0.72 ± 0.20 | 140 | 0.35 | 0.75 | 0.60 | 27 |
| ■ |  |  | ■ | 0.58 ± 0.07 | 0.75 ± 0.07 | 0.82 ± 0.05 | 1,259 | 0.52 | 0.70 | 0.82 | 257 |
| ■ | ■ |  | ■ | 0.51 ± 0.16 | 0.75 ± 0.15 | 0.75 ± 0.14 | 351 | 0.54 | 0.77 | 0.77 | 95 |
| ■ |  |  |  | 0.42 ± 0.04 | 0.78 ± 0.06 | 0.64 ± 0.06 | 1,259 | 0.32 | 0.68 | 0.64 | 257 |
| ■ | ■ |  |  | 0.39 ± 0.17 | 0.74 ± 0.15 | 0.65 ± 0.20 | 351 | 0.34 | 0.72 | 0.62 | 95 |

Two models showed consistent performance between the training and independent testing datasets: one combining GPS-derived predictors with environmental metrics and another combining GPS and wet-dry derived predictors with environmental metrics. In both models, sensitivity and specificity ranged from 0.70 to 0.82 across training and testing datasets (Table 1). We selected the model combining 11 GPS- derived movement metrics with environmental predictors, for subsequent analyses because it could also be applied to the dataset of birds equipped with radar detectors. In this model, environmental metrics ranked among the most important predictors (Fig. 2A2). Seabirds were more likely to attend vessels closer to the coast and within approximately 200 km of the colony (Fig. 2B2). Vessel-attendance was also more likely when sun azimuth ranged from −2 to 1 degrees, corresponding to early morning conditions (Fig. 2B2). Among GPS-derived movement metrics, vessel-attendance was associated with faster movement and greater maximum distances travelled within a given window (Fig. 2B2).

### 3.3 Discrimination between attendance at fishing and nonfishing vessels

After applying variable selection with the MUVR algorithhm, the models discriminating fishing from nonfishing vessel-attendance segments retained only a small subset of the original predictor variables. Between zero and four of the 15 GPS metrics were retained, together with zero to two of the seven wet-dry metrics, two to four of the 28 accelerometer metrics, and none of the seven pressure-derived metrics (Appendix S11).

Although several models achieved high sensitivity for identifying attendance at fishing vessels in the training dataset, specificity remained low, reaching a maximum of only 0.58 (Table 2). This indicates that more than 40% of nonfishing-vessel attendance segments were incorrectly classified as fishing-vessel attendance, showing limited discrimination between the two vessel types.

**Table 2.** Performance metrics of Random Forest models discriminating vessel-attendance at fishing from nonfishing vessels. Squares indicate the predictor groups included in each model. Models are ordered according to Youden’s J statistic. N indicates the number of trajectory segments in the training and testing datasets.

|  |  |  |  | Training |  |  |  | Testing |  |  |  |
|---|---|---|---|---|---|---|---|---|---|---|---|
| GPS | Wet-dry | Accel. | Envir. | Youden's J | Sensitivity | Specificity | N | Youden's J | Sensitivity | Specificity | N |
| ■ | ■ |  | ■ | 0.62 ± 0.24 | 0.98 ± 0.05 | 0.58 ± 0.30 | 172 | 0.22 | 0.92 | 0.30 | 47 |
| ■ |  | ■ |  | 0.54 ± 0.47 | 0.94 ± 0.08 | 0.50 ± 0.47 | 70 | 0.00 | 1.00 | 0.00 | 12 |
| ■ |  |  | ■ | 0.49 ± 0.13 | 0.95 ± 0.06 | 0.54 ± 0.11 | 633 | 0.45 | 0.97 | 0.48 | 131 |
| ■ | ■ |  |  | 0.37 ± 0.20 | 0.93 ± 0.08 | 0.40 ± 0.23 | 172 | 0.12 | 0.92 | 0.20 | 47 |
| ■ | ■ | ■ |  | 0.31 ± 0.27 | 0.64 ± 0.32 | 0.50 ± 0.50 | 31 | 0.00 | 1.00 | 0.00 | 14 |
| ■ | ■ | ■ | ■ | 0.24 ± 0.52 | 0.90 ± 0.09 | 0.20 ± 0.45 | 31 | 0.00 | 1.00 | 0.00 | 14 |
| ■ |  | ■ | ■ | 0.22 ± 0.44 | 0.98 ± 0.05 | 0.29 ± 0.40 | 70 | 0.25 | 1.00 | 0.25 | 12 |
| ■ |  |  |  | 0.14 ± 0.07 | 0.95 ± 0.04 | 0.19 ± 0.06 | 633 | 0.02 | 0.90 | 0.12 | 131 |

### 3.2 Agreement between vessel-attendance predictions and radar detections

To predict vessel-attendance along complete seabird trajectories, we applied the RF model combining GPS-derived movement and environmental predictors (Fig. 2A2–B2; Table 1). Predicted vessel-attendance probabilities were assigned to each GPS location along complete foraging trips, providing a continuous representation of the predicted probability of vessel-attendance (Fig. 3B). At the 1-h scale, the model discriminated periods with and without AIS-derived vessel-attendance with a balanced AUC of 0.85 (95% CI = 0.79–0.90).

**Figure 3.**
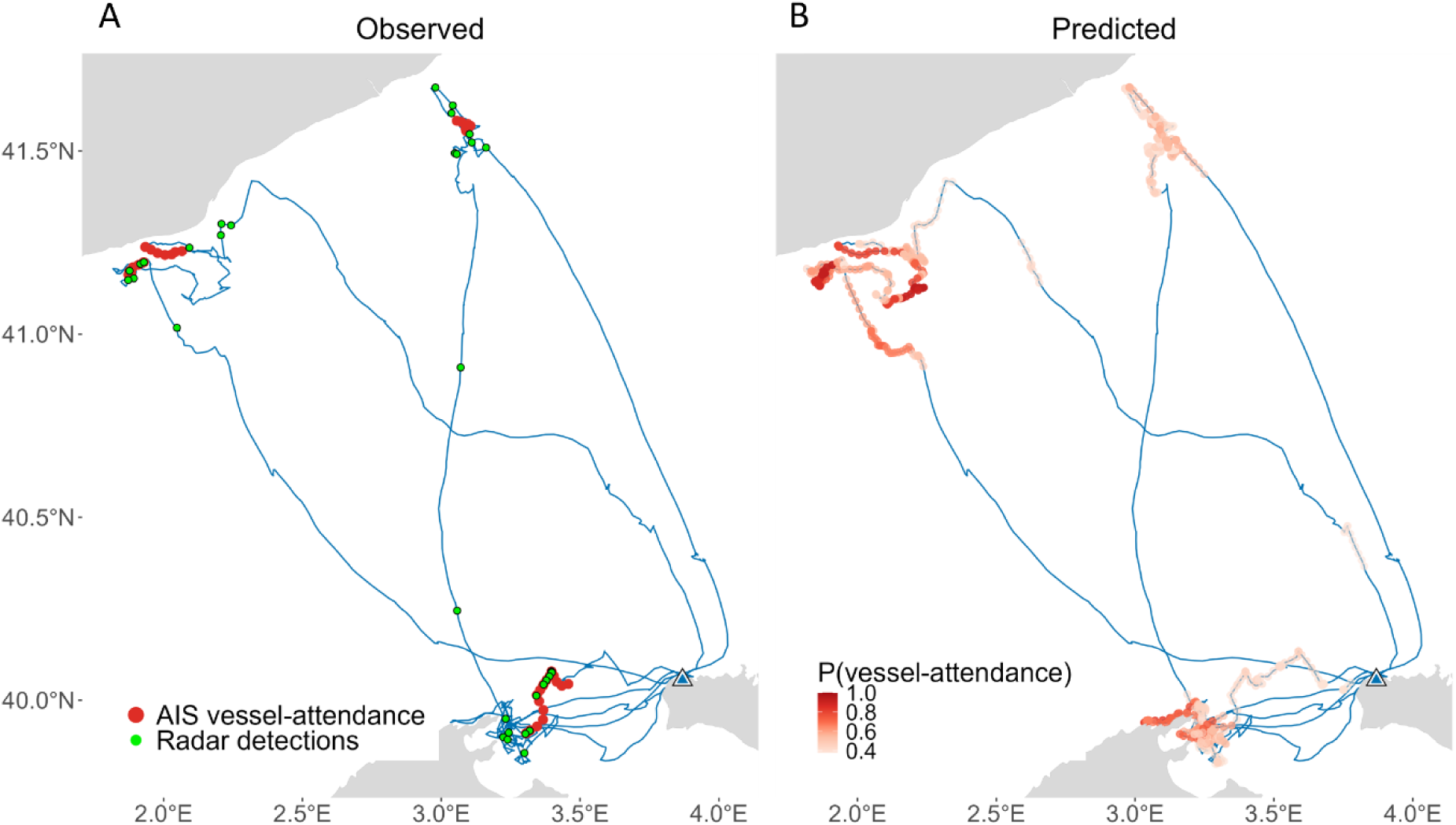
Example reconstruction of vessel-attendance along four Scopoli’s shearwater foraging trips. The left panel shows observed vessel-attendance locations identified from the spatiotemporal alignment of seabird GPS trajectories with AIS data (red) together with independent radar detections recorded by bird-borne radar detectors (green). The right panel shows continuous model support for vessel-attendance at each GPS location, derived from the Random Forest model combining GPS-derived movement and environmental predictors, with darker colours indicating stronger model support.

We then examined agreement at the level of individual events by converting predicted probabilities into binary classifications and grouping consecutive radar detections into events. At this fine scale, many predicted events were not supported by independent vessel information. Across complete foraging trips, we independently identified 227 predicted vessel-attendance events, 24 AIS-derived vessel-attendance events and 47 radar-detected vessel-presence events (Fig. 4; Appendix S12). After grouping temporally overlapping events, 39 of the 227 predicted events (17%) coincided with AIS or radar detections, whereas 188 (83%) had no corresponding detection from either source. Of these 39 events, 19 had no corresponding AIS-derived vessel attendance but coincided with radar detections of nearby vessels. Conversely, 16 groups containing radar detections did not overlap with any predicted vessel-attendance event.

**Figure 4.**
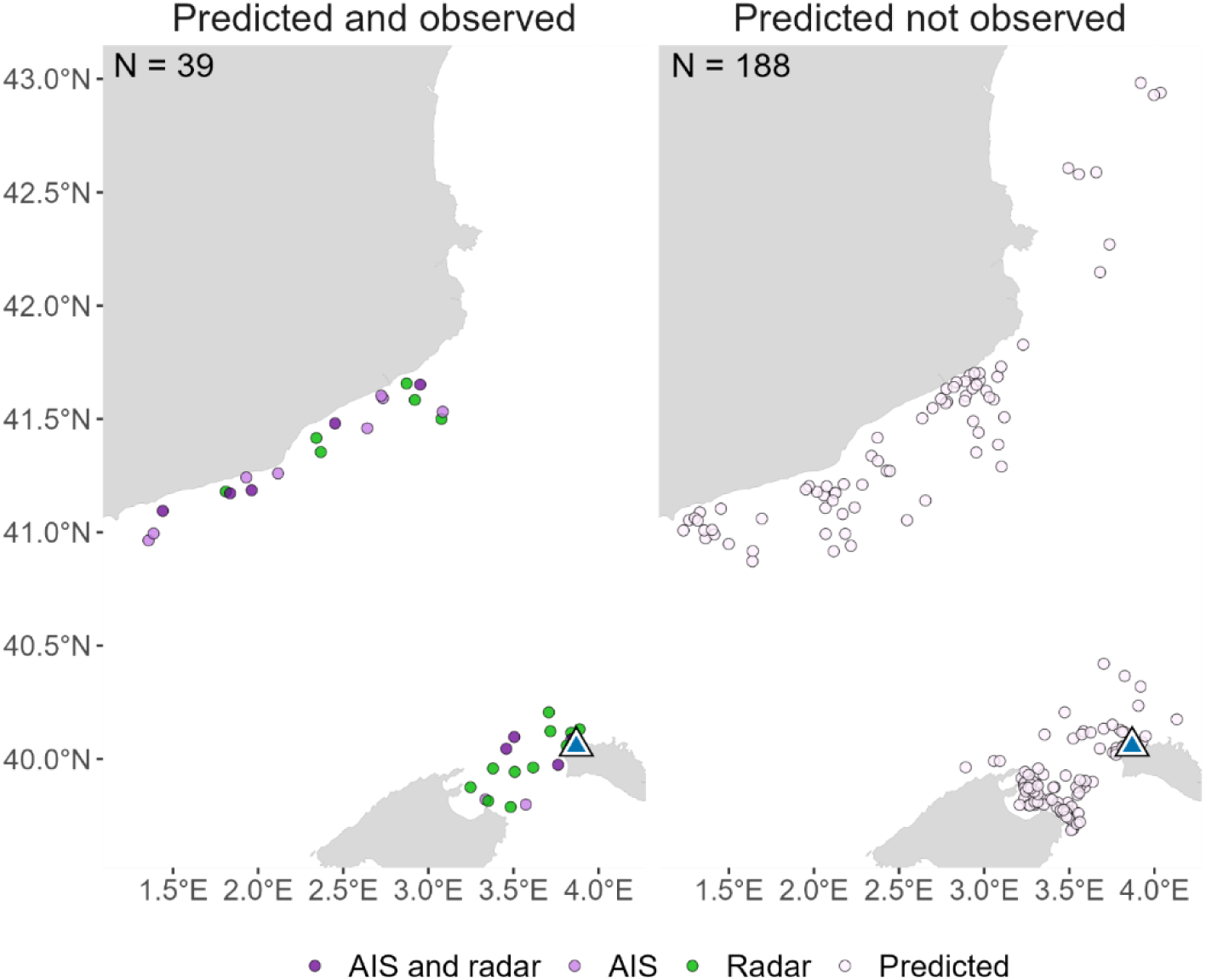
Comparison of all predicted vessel-attendance events with independent vessel-detection data. The left panel shows predicted events overlapping AIS derived vessel-attendance events, bird-borne radar detections, or both. The right panel shows predicted events lacking independent support from either AIS or radar detections. *N* indicates the number of predicted events in each category.

## 4 DISCUSSION

Using a comprehensive biologging dataset from shearwaters in an area of high maritime traffic, we show that vessel-attendance can be inferred from seabird movement and environmental information derived from bird-borne data, without direct access to vessel-tracking data. However, attendance at fishing and nonfishing vessels could not be reliably distinguished. Movement information alone did not generalise consistently across individuals, whereas incorporating environmental context improved model generalisation. Across complete foraging trips, the model combining GPS-derived movement and environmental predictors discriminated well between periods with and without AIS-derived vessel attendance at the hourly scale, but was less reliable at identifying individual events, many of which lacked AIS or radar support. Nevertheless, some predictions without AIS support coincided with nearby vessels detected by radar. Together, these findings highlight the potential of seabird biologging to help address gaps in estimating bycatch risk and complement existing vessel-tracking systems.

We found that a GPS-only deployment, when combined with a few environmental variables, could provide consistent predictions of vessel-attendance. Time of day, distance to the colony and distance to the coast substantially improved model performance, and the resulting patterns, such as higher attendance near continental shelves and during early morning hours, were consistent with previous studies on Scopoli’s shearwaters (Cianchetti-Benedetti et al., 2018; Genovart et al., 2018; Reyes-González et al., 2021; Soriano-Redondo et al., 2016). These environmental predictors are likely to capture, at least in part, regional patterns of vessel activity as well as species and region-specific patterns of seabird behaviour, and their importance may therefore reduce the transferability of our approach to other regions without local recalibration. Nevertheless, our findings support the potential application of this approach to larger existing GPS datasets to investigate seabird-vessel interactions across the Mediterranean.

When applied across complete foraging trips, the model discriminated periods with and without vessel-attendance at the 1-hour scale (AUC = 0.85). This suggests that the approach may be useful for identifying spatio-temporal patterns of vessel-attendance, particularly where vessel-tracking information is incomplete or unavailable. Such information could, for example, be useful in future demographic studies examining the consequences of seabird-vessel interactions. However, examining predictions at the level of individual events revealed substantially greater uncertainty. Predictions recovered most AIS-derived events and approximately two-thirds of the retained radar events, but only a relatively small proportion of all predicted events was supported by either source. Some predicted events without AIS support nevertheless coincided with radar detections of nearby vessels, providing evidence of vessel presence that was missing from AIS records. These events may have involved vessels intentionally not broadcasting AIS, although this appears uncommon in the study area (Navarro-Herrero et al., 2024) and has been associated with concealing fishing grounds (Ferrà et al., 2020; Welch et al., 2022), or recreational and artisanal vessels exempt from AIS requirements (Robards et al., 2016). Conversely, predictions unsupported by either AIS or radar may reflect model misclassification, conditions poorly represented during training, or interactions with vessels not detected by either system. Such predictions occurred in coastal areas potentially used by artisanal fisheries and recreational vessels, as well as further offshore, where terrestrial AIS coverage may be poorer. Distinguishing among these possibilities remains challenging when vessel-tracking coverage is incomplete, particularly for small-scale fisheries. This is especially relevant in the Western Mediterranean, where small-scale fisheries account for a substantial proportion of seabird bycatch (Cortés et al., 2017). The forthcoming implementation of mandatory vessel-tracking systems for European small-scale fisheries (Mendo et al., 2026) will provide an opportunity to evaluate this approach against a more complete representation of vessel activity.

Unlike recent studies on albatrosses, we were unable to distinguish attendance at fishing and nonfishing vessels from bird-borne data alone (Le Scornec et al., 2025; Rutter et al., 2025). Whereas albatrosses in those studies interacted almost exclusively with fishing vessels, more than one-third of vessel-attendance events in our study involved nonfishing vessels. These results suggest that identifying the presence of vessels and identifying the type of vessel represent two distinct prediction problems. The ability to discriminate between vessel types may also depend on the spatial context in which interactions occur. In regions where fishing and nonfishing vessels operate in more spatially distinct areas, spatial information associated with seabird movements could potentially improve vessel-type classification. In contrast, the spatial overlap among vessel types in our study area may have contributed to the limited discrimination observed here. In addition, in our study, attraction and following behaviour were grouped into a single vessel-attendance category (March et al., 2026), potentially obscuring behavioural differences associated with vessel type. Future studies could explore whether focusing specifically on following events improves discrimination between fishing and nonfishing vessels, although the number of such events in our dataset was limited. Improving our ability to discriminate fishing from nonfishing vessels should therefore be a priority, particularly in the Mediterranean, where fisheries bycatch remains one of the principal threats to Scopoli’s shearwater populations (Genovart et al., 2017; BirdLife International, 2024).

On the other hand, the relatively frequent attendance at nonfishing vessels indicates that maritime traffic itself can shape shearwater behaviour. Although our study considered attraction towards vessels, attraction is unlikely to be the only behavioural response to increasing vessel traffic, since avoidance responses have also been documented in several seabird species (Fliessbach et al., 2019). Extending individual-tracking approaches to identify separately attraction, following and avoidance could substantially improve our understanding of how maritime traffic influences habitat use and movement decisions of seabirds, and ultimately exposure to human activities.

Although movement-only models did not generalise consistently across individuals, the multi-sensor model combining GPS, wet-dry and accelerometer data achieved the highest predictive performance during training. GPS- and accelerometer-derived variables contributed most to predictions, followed by wet-dry data, whereas pressure-derived variables contributed little and were excluded during feature selection. However, the relatively small number of individuals equipped with multiple sensors limits our ability to determine whether the additional behavioural information provided by these sensors improves predictions across individuals. Larger multi-sensor datasets will therefore be needed to assess whether their higher performance during training translates into improved model generalisation and transferability. In this context, incorporating animal-borne cameras in future studies could also provide valuable opportunities for independent validation of vessel-attendance predictions.

Overall, our study demonstrates that biologging data can be used not only to understand seabird behaviour but also to reconstruct patterns of vessel activity at sea. Using only bird-borne tags, this approach can identify periods of potential vessel attendance when vessel-tracking data are incomplete or unavailable, and help revealing gaps in vessel-tracking coverage. By complementing existing vessel-tracking systems, these insights could improve assessments of fisheries pressure on seabirds, strengthen monitoring of poorly tracked fleets, and contribute to managing unreported vessel activities, including illegal practices, thereby supporting more effective conservation strategies to mitigate the impacts of fishing.

## Supporting information

Supplementary File 1 Reproducible R workflow

## Acknowledgements

Study funded by Spanish Government and EU FEDER (CGL2016-78530-R, PID2020-117155GB-I00/AEI/10.13039/501100011033 and PID2023-146782OB-I00/MICIU/AEI/10.13039/501100011033), PLEAMAR from *Fundación Biodiversidad* (2019/1423). Leia Navarro-Herrero was supported by Pleamar (2017/2349), BES-2017-079874 (CGL 2016-78530-R), “Pla GenT” program of the Generalitat Valenciana (CIDEGENT/2021/058) and “REDUCE” (EU Horizon Europe 2024, HORIZON-CL6-2023-BIODIV-01). We thank Sarah Saldanha for her advice in processing accelerometer and pressure sensor data, and Zuzana Zajková in processing wet-dry sensor data. David March and Jacob González-Solís acknowledge support from the Generalitat Valenciana through the “Pla GenT” program (CIDEGENT/2021/058) and from ICREA Acadèmia programme, respectively. We acknowledge support from the University of Exeter’s Advanced Research Computing facilities at Penryn in carrying out this work. We thank the Govern de les Illes Balears for permits and Andrea Soriano, Laura Zango, Fernanda Pereira, Montserrat Vanerio and many undergrad and master students for their help with the fieldwork.

## Author’s contribution

Leia Navarro Herrero: Software, validation, formal analysis, investigation, data curation, formal analysis, investigation, writing – original draft, visualization; David March: Conceptualization, methodology, software, validation, formal analysis, investigation, resources, data curation, writing – reviewed and editing, supervision, funding acquisition; Frederic Bartumeus: Conceptualization, methodology, writing – reviewed and editing; Jacob González-Solis: Conceptualization, resources, writing – reviewed and editing, supervision, project administration, funding acquisition.

## Data sources

Seabird tracking data can be downloaded upon request from the BirdLife International Seabird Tracking Database (https://www.seabirdtracking.org/). Vessel metadata used to identify fishing vessels can be freely downloaded from Global Fishing Watch (https://globalfishingwatch.org/) and the EU Fleet Register (https://webgate.ec.europa.eu/fleet-europa/search_en). Raw AIS data are available from the Balearic Islands Coastal Observing and Forecasting System (SOCIB) and EXMILE SOLUTIONS LTD and provided to the Cavanilles Institute of Biodiversity and Evolutionary Biology in the context of a data sharing agreement with SOCIB, accessed 2024-02-05.

## Statement on inclusion

Our study on seabird-fishing interactions in the NW Mediterranean Sea was conducted by local scientists, ensuring alignment with regional conservation priorities. This approach, leveraging bird-borne sensors, not only enhances local seabird conservation efforts but also offers a valuable method for studying similar interactions in other regions with complex maritime traffic and/or limited vessel tracking data.

