## Supplementary figures and images for "Predicting seabird-vessel interactions from bird-borne data alone"

### 6215396_20190704_221003_20190715_114503_calibrated.png

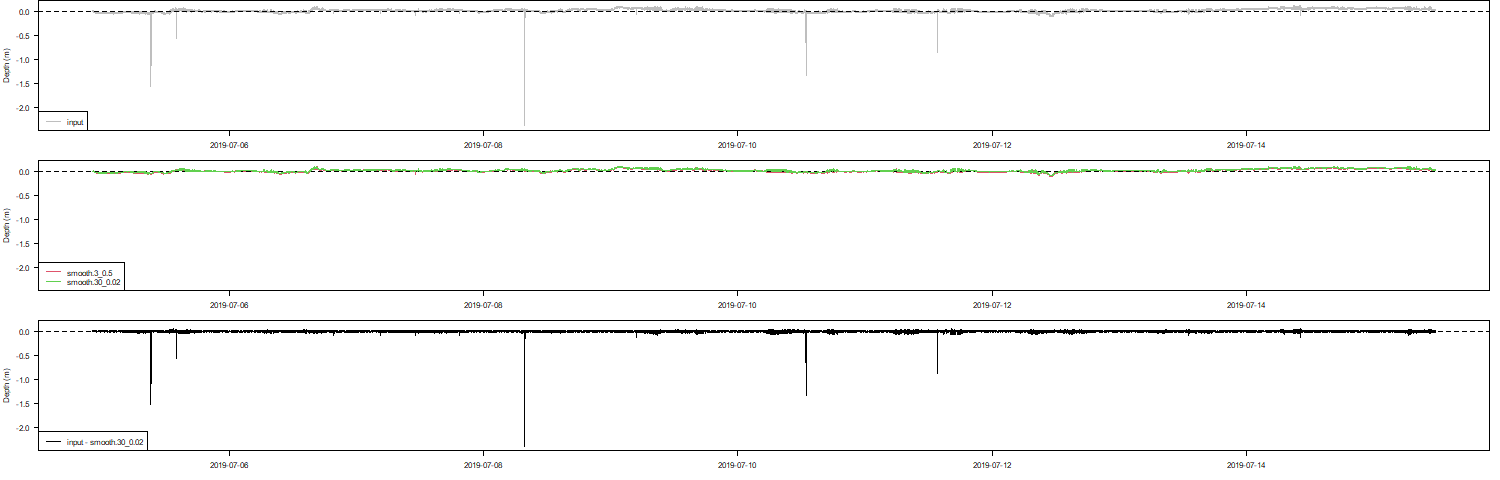

### 6215396_20190704_221003_20190715_114503_raw.png

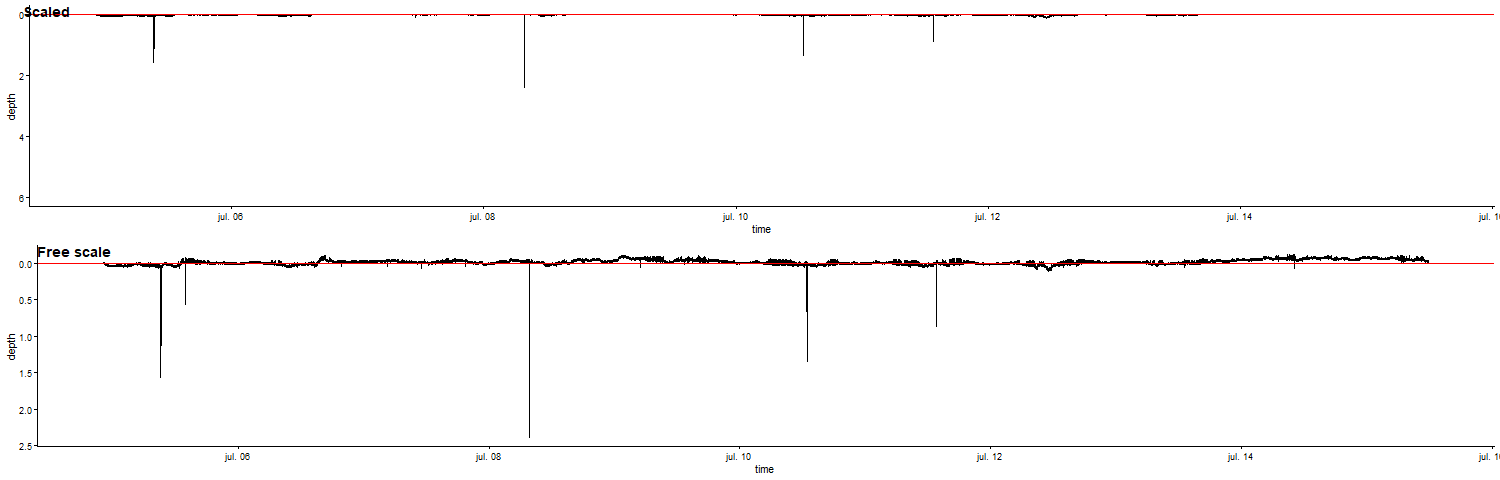

### duration_VesselSegments.png

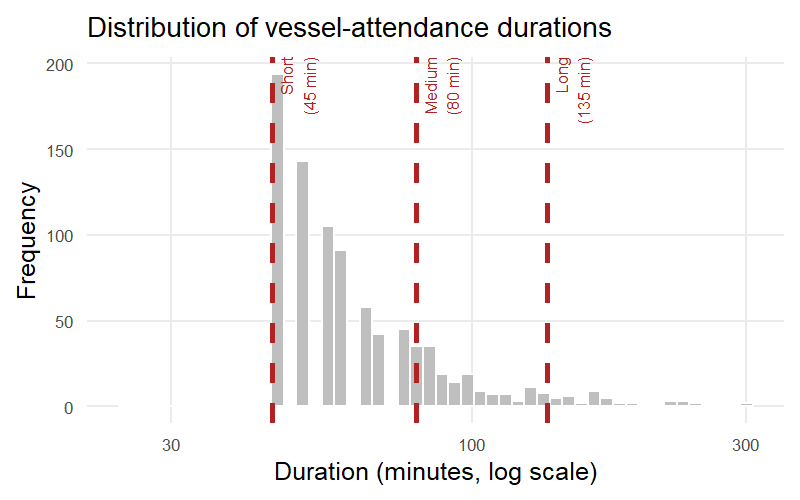

### min_GPS1_GLS0_ACC0_PRESS0_ST0_ClassA_Imp.png

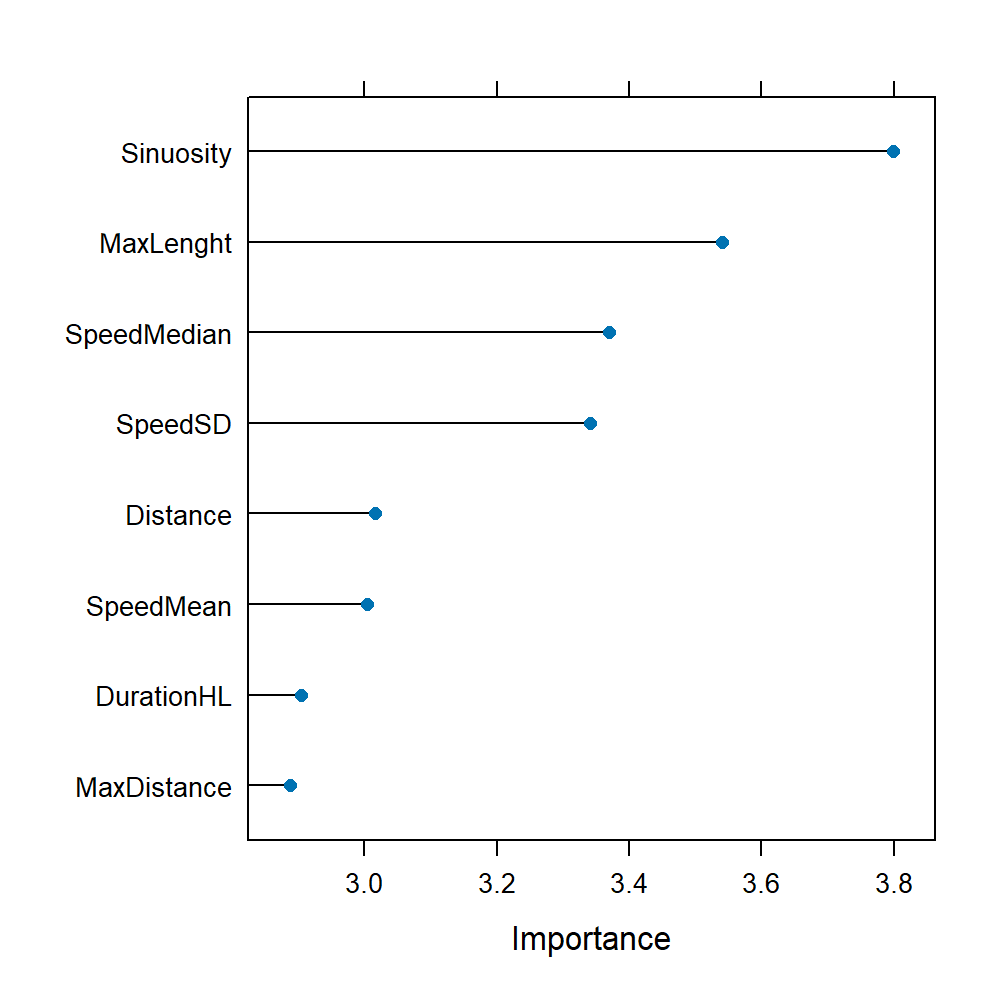

### min_GPS1_GLS0_ACC0_PRESS0_ST0_ClassA_peff.png

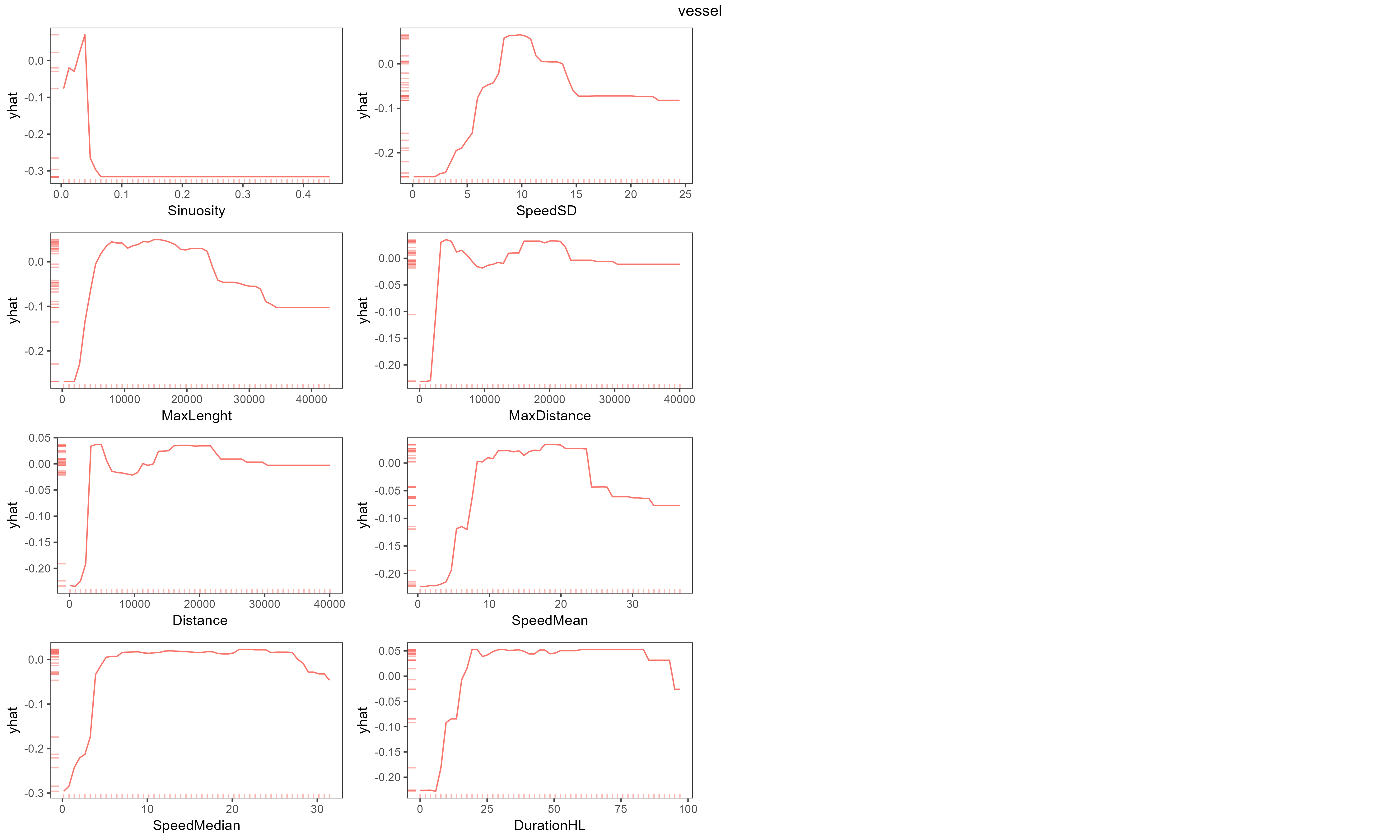
