## Supplementary File 2 Appendices for "Predicting seabird-vessel interactions from bird-borne data alone"

List of appendices:

**Appendix S1**: Methods - Procedure for generating pseudo-absence segments

**Appendix S2**: Table - Metrics calculated from GPS sensor.

**Appendix S3**: Table - Metrics calculated from wet-dry sensor.

**Appendix S4:** Methods - Data preprocessing: accelerometer and pressure data.

**Appendix S5**: Table - Metrics calculated from accelerometer sensor.

**Appendix S6**: Table - Metrics calculated from pressure sensor.

**Appendix S7**: Table - Environmental metrics calculated from GPS sensor.

**Appendix S8:** Figure - Distribution of observed vessel-attendance durations.
**Appendix S9:** Methods - Assessment of temporal autocorrelation to determine the temporal scale for model evaluation.
**Appendix S10**: Figure - Distribution of behaviour percentages within vessel-attendance segments.
**Appendix S11**: Table - Minimal Variable Selection Random Forest-based Algorithm (MUVR).
**Appendix S12:** Figure - Overlap among predicted, radar and AIS vessel-attendance events.

**Supplementary References**

**Appendix S1**. Procedure for generating pseudo-absence segments.

To assess whether seabird movement differed between vessel attendance and natural movement, we used trajectory segments as the units of analysis. Vessel-attendance segments were treated as presences and comparable natural-movement segments as pseudo-absences.

Because seabirds may detect vessels before approaching them closely we incorporated the 30 minutes preceding each vessel-attendance event into the analysis (Corbeau et al., 2019; Votier et al., 2010; March et al., 2026). This procedure defined what we hereafter refer to as a *vessel-attendance segment*. For each vessel-attendance segment, we then generated a pseudo-absence by randomly selecting a comparable trajectory segment in which no vessel-attendance event occurred. These segments are hereafter referred to as *natural-movement segments*. To ensure comparability, pseudo-absence segments were required to have similar attributes to the corresponding vessel-attendance segments, specifically the same duration and a comparable behavioural pattern. Behavioural states were assigned for each individual trajectory using the Expectation-Maximisation binary Clustering algorithm (EMbC; Garriga et al., 2016). This algorithm classifies movement fixes based on speed and turning angle into four behavioural modes: intensive search (low speed and high turning angles), extensive search (high speed and high turning angles), resting (low speed and low turning angles), and relocating (high speed and low turning angles). Following previous studies, intensive and extensive search behaviours were grouped as foraging behaviour (Reyes-González et al., 2021).

We then classified each vessel-attendance segment according to its predominant behavioural pattern (foraging, resting, or relocating). If more than 80% of the fixes within a segment were classified as relocating or resting, the segment was assigned to that behavioural category; otherwise, it was classified as foraging.

**Appendix S2**. Metrics calculated from GPS. Code is available in Data availability statement.

| Metric | Units | Definition |
| --- | --- | --- |
| Length | m | Total distance travelled (trajr R Package McLean & Skowron Volponi, 2018) |
| Distance | m | Net distance between the start and the end (trajr R Package McLean & Skowron Volponi, 2018) |
| Max. length | m | Total distance travelled between the start and the farthest point from the start (trajr R package McLean & Skowron Volponi, 2018) |
| Max. distance | m | Net distance between the start and the farthest fix from the start (trajr R package McLean & Skowron Volponi, 2018) |
| Max. step length | m | Distance of the longest step (two consecutive fixes) in terms of distance (trajr R package McLean & Skowron Volponi, 2018) |
| Sinuosity | 0 - 1 | Tortuosity measurement by Benhamou, S. (2004), where a value of 0 indicates a straight path and a value of 1 indicates a highly curved path (trajr R package McLean & Skowron Volponi, 2018) |
| Straightness | 0 - 1 | Opposite concept of tortuosity. It is calculated as the ratio between distance and length, where a value of 0 indicates infinitely tortuous and a value of 1 indicates a straight line (trajr R package McLean & Skowron Volponi, 2018) |
| Speed | km/h | Distance travelled divided by time taken (move R Package^,^ Kranstauber et al., 2018 ). Speed values are summarized with the mean, SD, median and maximum speed value. |
| EMbC state duration | min | Total duration in each EMbC state (HH, HL, LL, LH) based on estimates of speed and turning angles (Garriga et al., 2016). "H" indicates high, "L" indicates low. The first letter corresponds to speed, and the second letter indicates turning angles. |

**Appendix S3**. Metrics calculated from wet-dry sensor. Code is available in Data availability statement.

| Metric | Units | Definition |
| --- | --- | --- |
| Rate of state transitions | N transitions/min | Number of state transitions divided by the total duration of the segment |
| Landings |  | Number of transitions from dry to wet states |
| Wet state duration | min | Total duration in the wet state |
| Max. wet state duration | min | Maximum duration in a sustained wet state |
| Dry state duration | min | Total duration in the dry state |
| Max. dry state duration | min | Maximum duration in a sustained dry state |
| Proportion dry state | 0 - 1 | Total duration in a dry state divided by the total duration of the segment |

**Appendix S4**. Data preprocessing: accelerometer and pressure data.

*Accelerometer data*

We processed accelerometer data sampled at 25 Hz (25 observations per second) to derive key movement metrics at the segment level. To obtain segment-level summaries, we first computed the mean and standard deviation (SD) for each metric (roll and pitch metrics, dynamic acceleration along the X, Y, and Z axes, as well as ODBA and VeDBA metrics; see Supplementary Table S5 for detailed calculations) at one-second intervals, by averaging across the 25 observations per second, aligning with the recommendations (Cianchetti-Benedetti et al., 2017; Collins et al., 2015). This step was necessary to reduce noise and account for rapid fluctuations in behavior that occur at very short timescales. But the final data used in our analysis consisted of aggregated metrics at the segment level. For each metric, we derived:

- The mean of all one-second means, to represent the average behavior over the entire segment.
- The mean of all one-second SDs, to capture the average variability within each second across the segment.
- The SD of all one-second means, indicating the overall variation in behavior across the segment.
- The SD of all one-second SDs, representing the variability in the short-term fluctuations within the segment.

By calculating these metrics, we effectively summarised both the typical movement patterns and the variability in those patterns over time in each segment. Code is available in Data availability statement.

*Pressure data*

We determined the pressure mode (Pm) for each trip and then derived the depth (D, in meters) through the equation: D = 0.01 × (Pm − Pa), where a 1-millibar difference in pressure corresponds to a 0.01-meter variation in depth, in accordance with a previous work (Saldanha et al., 2023). To account for surface drift, we applied the zero-offset correction with the ‘diveMove’ R package R (Luque & Fried, 2011). Given the sensitivity of the pressure sensors our analysis only considered dives of 0.5 meters or deeper. Code is available in Data availability statement.

**Supplementary Table S5**. Metrics calculated from accelerometer sensor. Code is available in Data availability statement.

| Metric^1^ | Equation | Definition |
| --- | --- | --- |
| Surge |  | X acceleration (g). Reflects forward and backwards motion of the longitudinal X axis |
| Sway |  | Y acceleration (g). Reflects side to side motion (horizontal movement) of the longitudinal Y axis |
| Heave |  | Z acceleration (g). Reflects upward and downward motion (vertical movement) of the vertical Z axis |
| Pitch | $Arctan\sqrt{\frac{X}{Y^{2}+Z^{2}}}*\frac{180}{\pi}$ | Reflects rotation from head to tail. |
| Roll | $Arctan\sqrt{\frac{Y}{X^{2}+Z^{2}}}*\frac{180}{\pi}$ | Reflects rotation from side to side |
| ODBA | $a_{X}+ a_{Y}+ a_{Z}$ | Overall dynamic body acceleration reflects overall movement intensity. It is calculated as the sum of absolute dynamic accelerations (a) of the three axes (X, Y, Z). Dynamic acceleration is calculated as acceleration on the axis minus the static acceleration on the same axis. |
| VeDBA | $\sqrt{{(a}_{X}^{2} {+a}_{Y}^{2}+a_{Z}^{2}}$ | Vectorial dynamic body acceleration reflects overall movement direction and magnitude. It is calculated as the squared root of the sum of squared dynamic accelerations (a, see ODBA) of the three axes (X, Y, Z) |

^1^ Each metric was summarized at the segment level by calculating the mean and standard deviation (SD) from the means and SDs obtained at one-second intervals (see Appendix S5 for more detail).

**Appendix S6**. Metrics calculated from pressure sensor. Code is available in Data availability statement.

| Metric | Units | Definition |
| --- | --- | --- |
| Dives |  | Number of dives (diveMove R Package Luque & Fried, 2011) |
| Max. depth | m | Maximum depth (diveMove R package Luque & Fried, 2011) |
| Duration descending | min | Total duration descending on dives (diveMove R package Luque & Fried, 2011) |
| Duration on the bottom | min | Total duration on the bottom (diveMove R package Luque & Fried, 2011) |
| Duration ascending | min | Total duration ascending from dives (diveMove R package Luque & Fried, 2011) |
| Max. duration diving | min | Maximum duration of a dive (diveMove R package Luque & Fried, 2011) |
| Duration diving | min | Total duration of dives (diveMove R package Luque & Fried, 2011) |

**Appendix S7**. Environmental metrics calculated from GPS sensor**.** Code is available in Data availability statement.

| Metric | Units | Definition |
| --- | --- | --- |
| Altitude | ° | Sun angular height above horizon, 0° to 90°. High altitudes during daytime, low altitudes at sunrise and sunset (suncalc R Package Thieurmel, 2019) |
| Azimuth | ° | Sun's direction on the horizon, 0° to 360° circle. Morning east (90°), afternoon west (270°) (suncalc R Package Thieurmel, 2019) |
| Colony distance | km | Distance to breeding colony considering land barriers (raster R Package Hijmans, 2017) |
| Coast distance | km | Distance to closest coast considering land barriers (raster R Package Hijmans, 2017) |

**Appendix S8**. Distribution of observed vessel-attendance event durations. Red dashed lines indicate the three temporal window lengths selected using logarithmic spacing to span the central 90% of the observed duration distribution.

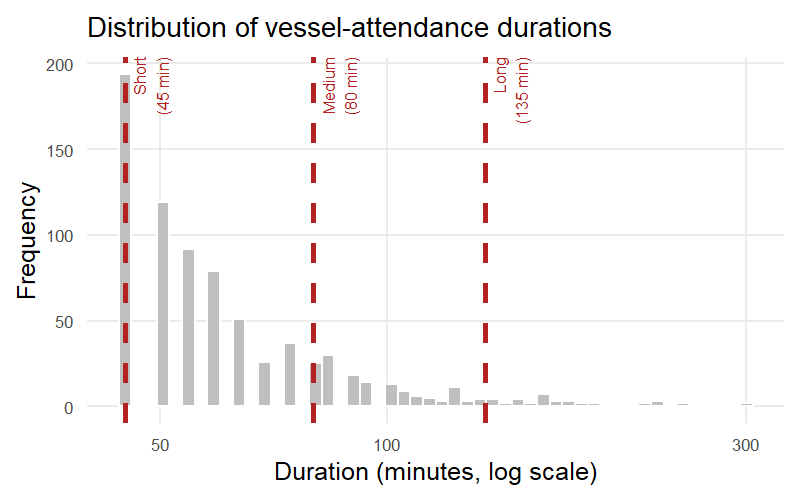

**Appendix S9.** Assessment of temporal autocorrelation to determine the temporal scale for model evaluation.

To select an appropriate temporal scale for model evaluation, we examined temporal autocorrelation in the observed vessel-attendance state (0/1) independently for each foraging trip using the acf function from the R *stats* package. As GPS locations were available at 5-minute intervals, we calculated autocorrelation at lags corresponding to 30 min, 1 hour and 2 hours.

Median autocorrelation across trips decreased from 0.53 at 30 minutes to 0.20 at 1 hour and −0.02 at 2 hours, indicating that temporal dependence was substantially reduced at 1 hour and negligible at 2 hours. Moreover, vessel-attendance events lasted on average 38.3 ± 33.8 minutes, suggesting that consecutive 30-minute intervals could frequently fall within the same event. We therefore selected 1-hour bin as a compromise between reducing temporal autocorrelation and retaining sufficient temporal resolution for model evaluation.

**Appendix S10.** Distribution of behavioural-state percentatges within vessel-attendance segments. Each observation within the boxplot represents behaviour percentage (Garriga et al., 2016), providing an overview of behaviour patterns across fishing and nonfishing vessels.

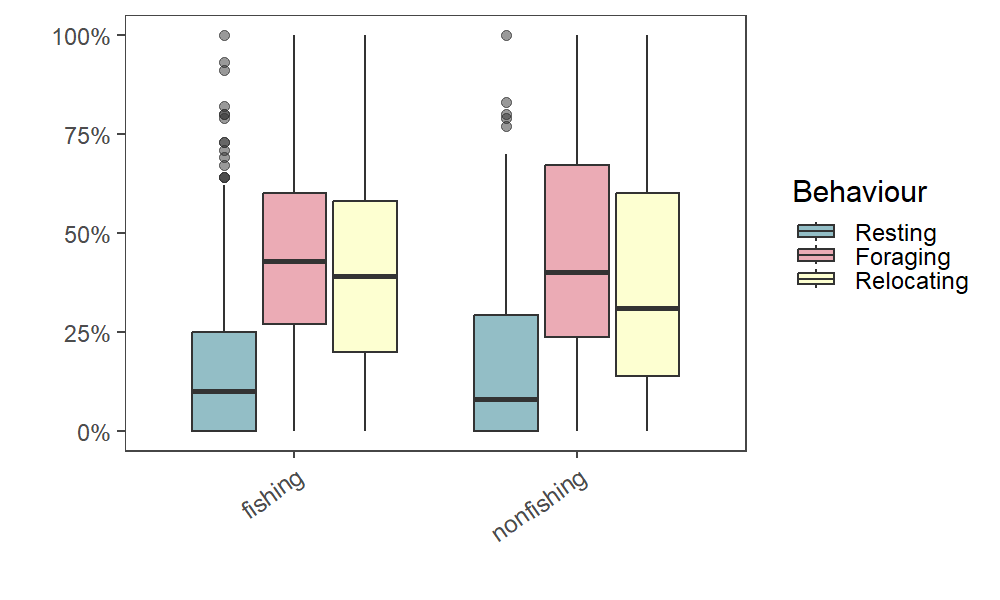

**Appendix S11*.*** Minimal Variable Selection Random Forest-based Algorithm (MUVR) for models comparing vessel-attendance and natural movement segments (A), and for comparing fishing and nonfishing vessel-attendance segments (B). Sensor selection results obtained from the Minimal Variable Selection Random Forest-based Algorithm (MUVR; Shi et al., 2019). Each row represents a model, indicating the number of variables of each sensors considered in the input ('i') and how many they were retained in the output ('o') after performing MUVR and the number of observations (N obs) included in each model, and the classification rate (CR, percentage of correctly classified observations obtained during the MUVR cross-validation procedure).

(A) vessel-attendance and natural movement segments

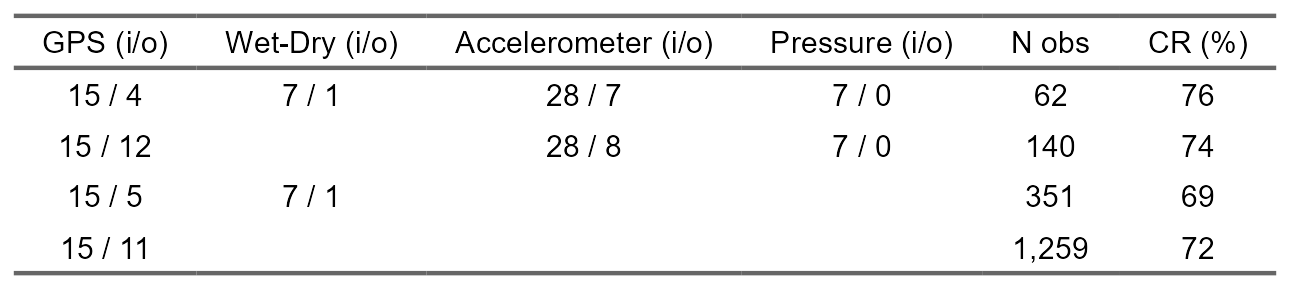

(B) fishing and nonfishing vessel-attendance segments

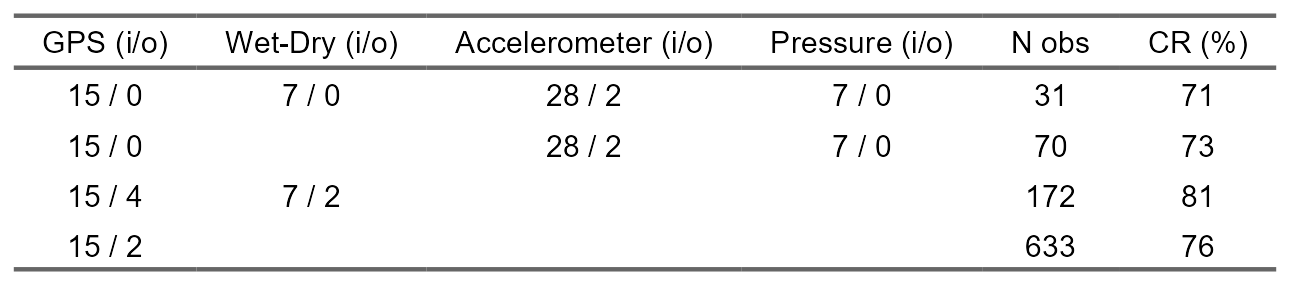

**Appendix S12***.* Overlap among predicted vessel-attendance events, radar detections and AIS vessel-attendance events. Consecutive detections separated by less than 30 min were grouped into a single event. Blue cells indicate the data source(s) contributing to each overlap category, and numbers on the right show the number of merged events in each category.

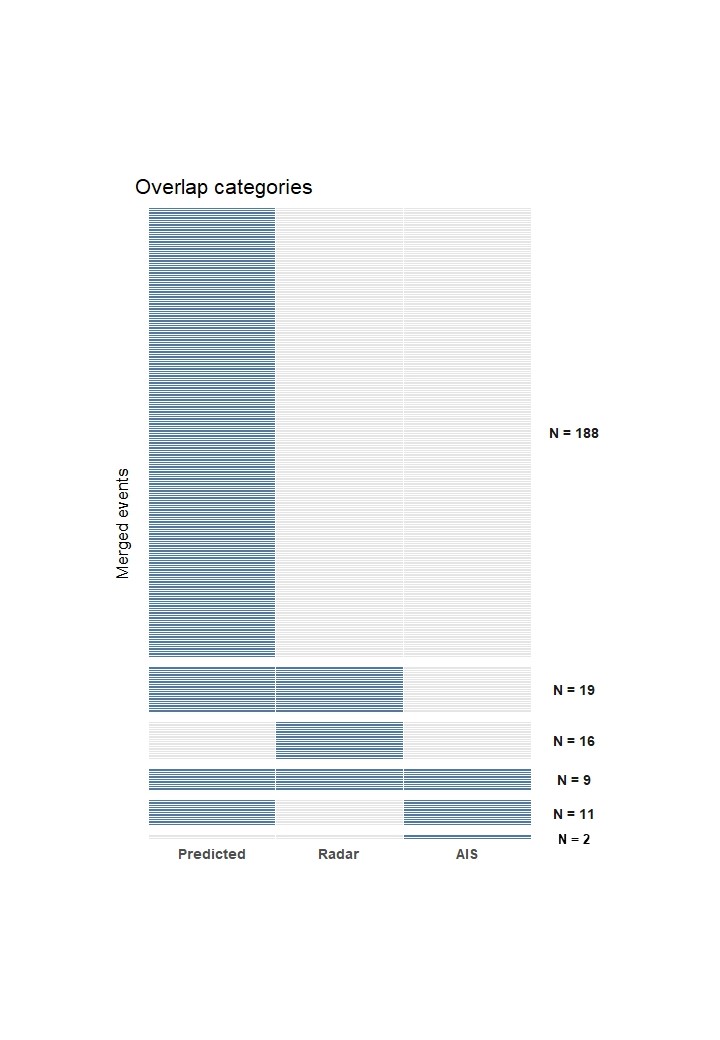
